# SCC*mec*Extractor: A tool for extracting Staphylococcal Cassette Chromosome elements from Whole Genome Sequences

**DOI:** 10.64898/2026.03.31.715619

**Authors:** Alison C. MacFadyen

## Abstract

Staphylococcal cassette chromosome (SCC) elements are mobile genetic elements that integrate at the *rlmH* gene and are predominantly responsible for methicillin resistance in staphylococci. Although SCC*mec* typing tools exist, none can extract the element sequence itself or explicitly classify SCC elements that lack methicillin resistance genes. Here we present SCC*mec*Extractor, a lightweight Python toolkit that identifies SCC element boundaries through degenerate attachment site (*att*) pattern matching, extracts complete elements from whole-genome assemblies and characterises their *mec* and *ccr* gene content. Benchmarking on 7,297 genomes spanning 70 species across *Staphylococcus* and *Mammaliicoccus* demonstrated 100% typing concordance with the `sccmec` tool^1^ on 1,454 *S. aureus* genomes. The tool extracted 1,562 SCC elements, from 1,454 *S. aureus*, 5,295 non-*aureus Staphylococcus* and 548 *Mammaliicoccus* genomes, achieving effective extraction rates (excluding assembly-limited genomes and those lacking valid *ccr* pairs) of 87.3% for *S. aureus*, 58.8% for non-*aureus Staphylococcus*, and 61.9% for *Mammaliicoccus*. Notably, 616 of the 1,562 extracted elements (39.4%) were non-*mec* SCC elements lacking methicillin resistance genes, a class of mobile element often overlooked. Non-*mec* SCC prevalence increased from 12.2% in *S. aureus* to 55.6% in non-*aureus Staphylococcus* and 76.0% in *Mammaliicoccus*, revealing a substantial reservoir of SCC diversity beyond methicillin resistance. SCC*mec*Extractor is freely available via PyPI, Docker and Singularity under an MIT licence.

**Impact Statement:** Staphylococcal cassette chromosome (SCC) elements are mobile genetic elements responsible for methicillin resistance in staphylococci and are central to methicillin resistant *Staphylococcus aureus* (MRSA) epidemiology. Existing tools focus on typing SCC*mec* from assemblies but cannot extract the element itself, limiting our ability to comprehensively monitor and examine these elements. SCC*mec*Extractor is a lightweight, portable tool that detects the attachment sites, required by SCC elements to integrate into the genome, extracts the SCC element, both *mec* gene carrying and not, and characterises their gene content. Applied across 7,297 genomes spanning two genera, we demonstrate that non-*mec* SCC elements are the dominant SCC class outside *S. aureus*, a finding enabled by systematic extraction and classification of SCC elements regardless of mec gene content. SCC*mec*Extractor provides the research community with an accessible, confidence-first approach (based on biology) to SCC element analysis across all staphylococci and mammaliicocci species.

**Data Summary:** The code for this pipeline is available at: https://github.com/AlisonMacFadyen/SCCmecExtractor, with a Docker image available at: https://hub.docker.com/repository/docker/alisonmacfadyen/sccmecextractor and PyPi package at: https://pypi.org/project/sccmecextractor/. All reference databases are bundled with the tool. Benchmarking genome accessions: 1,454 *S. aureus*, 5,295 non-aureus *Staphylococcus*, and 548 *Mammaliicoccus* genomes from NCBI. A complete list of genome accessions is provided as supplementary data (Supplementary Table S1). Extracted SCC elements can be obtained from Zenodo: 10.5281/zenodo.19355206

## Introduction

Methicillin-resistant *Staphylococcus aureus* (MRSA) is an important human and animal pathogen^2^, with MRSA being a major cause of human death occurring due to antimicrobial resistance worldwide^3^. Acquisition of *mec* genes, most commonly *mecA*, confers resistance to methicillin in *staphylococcus* species, by encoding an alternative penicillin-binding protein (PBP2a) with low affinity for beta-lactam antibiotics^4,5^. These *mec* genes are carried on staphylococcal cassette chromosome *mec* (SCC*mec*), a mobile genetic element that integrates site-specifically into the chromosome at the 3’ end of the *rlmH* gene (formerly *orfX*)^6,7^.

Integration and excision of SCC*mec* is achieved via element carrying large serine recombinases, CcrAB or CcrC, which identify specific DNA attachment sites (*att*)^8^. For integration, a pair of *att* sites, *attB* on the chromosome and *attSCC* on the circularised SCC*mec* element, are required. After integration, two *att* sites are generated flanking the SCC*mec* element: *attR* (*rlmH*-proximal) and *attL* (distally located)^8^. Each *att* site comprises two half-sites flanking a central GA dinucleotide where strand exchange occurs; these half-sites form degenerate inverted repeats^9^. The half-sites are shared between *att* sites: three are similar to one another and derive from the SCC*mec* element, while the fourth, overlapping the *rlmH* coding frame, is found within *attB* and *attL*^9^. All four full *att* sites contain a conserved 8 bp core motif (TATCATAA), including a central cytosine thought to be essential for recombination^10^. Our *attR* search patterns capture the region spanning 3 bp upstream to 12 bp downstream of this core motif, while *attL* patterns target the SCC-derived half-site upstream of the core, including the variable bridge region that reflects the diversity of integrated elements.

SCC*mec* carry two essential genetic components: the *mec* gene complex, comprising the *mec* gene and in some cases the regulatory elements (*mecR1*, *mecI*), with the other component being the *ccr* gene complex^11^, encoding the large serine recombinase (*ccrA*/*ccrB* pairs or *ccrC*)^12^. The remainder of the element consists of joining (J) regions that are varied and can be highly diverse^13^, these regions are also the basis of defining SCC*mec* subtypes^12^. The International Working Group on the Classification of Staphylococcal Cassette Chromosome Elements (IWG-SCC) classifies SCC*mec* types based on the combination of *ccr* gene complex type (determined by *ccr* allotype) and *mec* gene complex class (determined by regulatory genes and insertion sequences)^14^. To date, 15 SCC*mec* types (I-XV) have been defined for *S.* aureus^12,15,16^.

SCC elements are not limited to SCC*mec*, with staphylococci carrying a diversity of SCC elements which lack methicillin resistance genes. These non-*mec* SCC elements retain the hallmarks of SCC, the *att* site boundaries, *ccr* recombinases and integration at *rlmH* but carry a variety of different genes e.g. mercury resistance (SCC*Hg*), capsule biosynthesis (SCC*cap1*), etc^17–19^. Despite their potential roles in bacterial adaptation and as vehicles for horizontal gene transfer, non-*mec* SCC elements are largely overlooked because existing tools focus on methicillin resistance.

Several bioinformatic tools have been developed for SCC*mec* analysis. The `sccmec` tool^1^ uses a targets and regions BLAST approach to assign formal SCC*mec* types I-XV from assembly data. However, it does not detect *att* sites or extract element sequences and although it identifies the presence of *ccr* genes, it does not specifically name non-*mec* SCC elements. SCC*mec*Finder, developed by the Center for Genomic Epidemiology, combines BLAST and k-mer approaches for typing, but the standalone tool requires Python 2.7 and legacy BLAST, making it effectively non-functional; its web server is functional but limited to types I-XI^20^. Similar to `sccmec`, SCC*mec*Finder identifies the presence of *ccr* genes in the absence of *mec* and will state that the organism is most likely methicillin sensitive *S. aureus* but does not formally identify the element as a SCC. Finally, SCC*mec*_CLA performs extraction of SCC*mec* elements. This is achieved by searching the distal *att* site using the 8 bp core motif (TATCATA[AT]), with BLAST validation against known references, then assigns the start of the element as the 3’ region of *rlmH*, without explicit *att* site detection. Extracted cassettes are typed (I-XI) through gene synteny for *mec* class and *ccr* allotype assignment, which is supplemented with k-nearest-neighbour subtype prediction using Mash distances to 267 reference cassettes. Although a sophisticated approach, it requires all three core components (*rlmH*, *mecA*, *ccr*), therefore cannot detect non-*mec* SCC elements, is limited to *S. aureus* by its single-species *rlmH* reference, and has undocumented dependencies that prevent reproducible installation.

A gap therefore exists for a lightweight, cross-platform tool that can: (i) detect SCC element boundaries through explicit *att* site identification; (ii) extract complete element sequences; (iii) characterise gene content at the allotype level; (iv) detect non-*mec* SCC elements; and (v) operate across staphylococcal and mammaliicoccal species. SCC*mec*Extractor addresses this gap.

## Implementation

Integration of SCC elements requires two key elements, the presence of recombinase, to carry out the integration, and attachment sites for recognition of the recombination site. Therefore, a key approach of SCC*mec*Extractor is to identify those attachment, or *att* sites, then determine if the genes encoding the recombinases, *ccr* genes, are present between the *att* site location. Thus, incorporating our biological understanding of how these elements integrate to identify them and extract their sequences from whole genome sequence data. The primary functions of SCC*mec*Extractor are: `sccmec-locate-att`, `sccmec-extract, sccmec-type`, `sccmec-report` and `sccmec-pipeline`, Figure 1 outlines the workflow. Input files can either be FASTA/FNA only or FASTA/FNA with a GFF3 annotation file.

**Figure 1:**
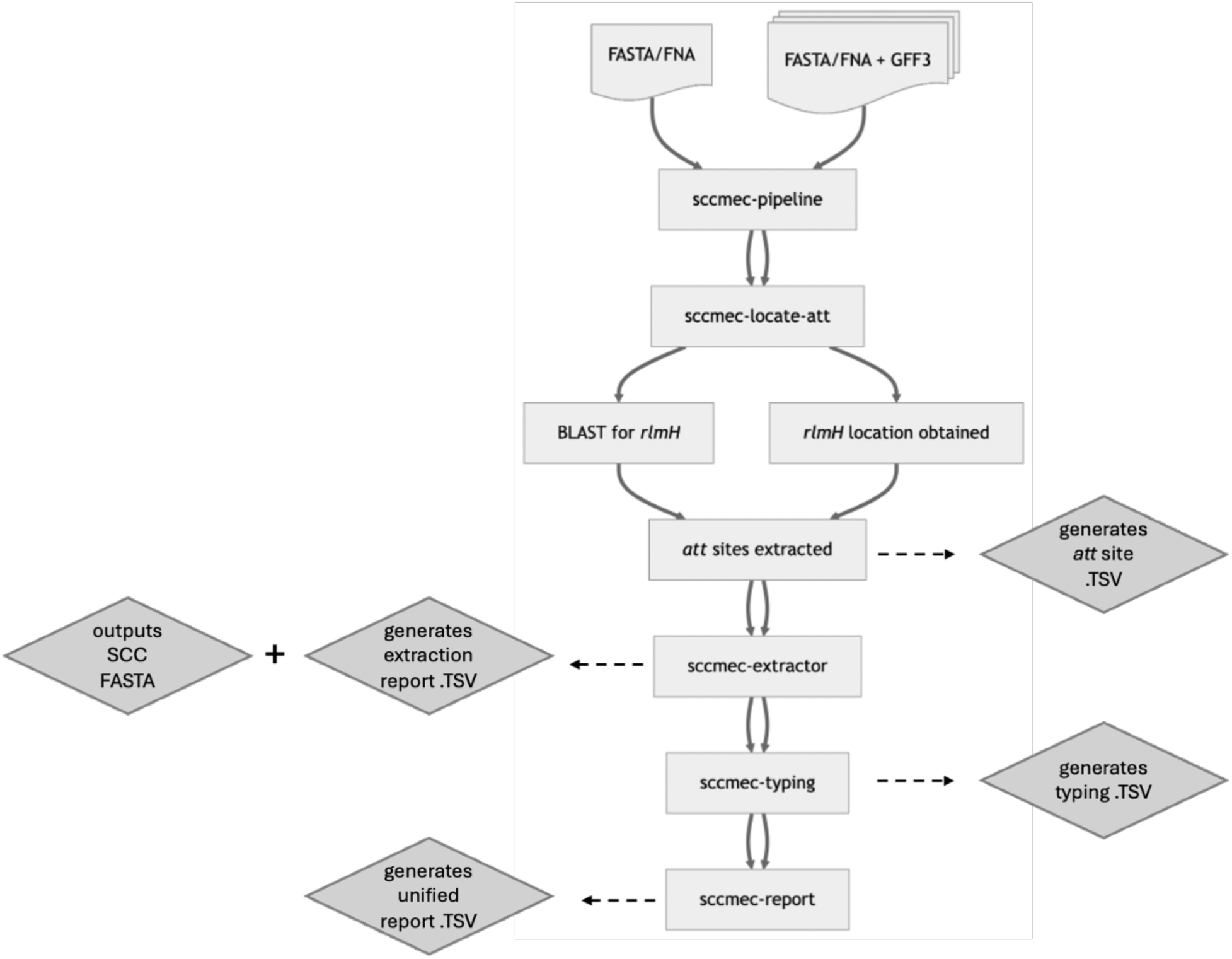
Schematic overview of the SCC*mec*Extractor pipeline showing the five command-line tools and their data flow. Input genomes (FASTA/FNA with optional GFF3) are processed through: (1) sccmec-locate-att for attachment site detection with *rlmH* validation; (2) `sccmec-extract` for SCC element extraction; (3) `sccmec-type` for BLAST-based gene-level typing of *mec* and *ccr* allotypes; (4) `sccmec-report` for merging extraction and typing results into a unified report. The master pipeline `sccmec-pipeline` orchestrates all steps in a single command. Arrows indicate data flow between modules; dashed arrows show the files/outputs produced by that step/tool.

**Attachment Site Detection -** sccmec-locate-att

As integration of SCC elements requires identification of *attB* and recombination with *attSCC*, resulting in the generation of *attR* and *attL* flanking the region, the first step in the workflow is to locate these *att* sites. Detection of these sites is achieved using 20 regular expression patterns, 4 forward and 4 reverse-complement for *attR* variants, with 6 forward and 6 reverse-complement patterns for *attL* variants, capturing species-specific bridge sequence diversity (Supplementary Table S2). All patterns were validated by cross-referencing detected *att* site coordinates against GFF3 gene annotations across all 7,297 benchmarking genomes, confirming that *attL* sites fall in intergenic regions while *attR* sites fall within *rlmH* as expected. Further to this, the majority of *att* sites contained the `GA` dinucleotide, located 6 bp upstream of the central 8 bp core motif, which is hypothesised to be where DNA strand exchange occurs during excision/integration^9^. The only exception was *attL14*, which carries a variant core motif (TATCCTAA) and an `AA` dinucleotide in place of `GA`. Despite this sequence divergence, *attL14* was retained as a valid pattern because it is exclusively associated with *ccrA2*/*ccrB2* at standard identity thresholds, paired with canonical *attR*, is restricted to *S. aureus* clinical isolates and consistently defines a distal SCC*mec* boundary flanked by IS21 elements, consistent with a genuine, if divergent, recombination product.

The variance within the patterns accounts for diversity across species. For example, the core *attR* pattern GC[AGCT]TATCA[TC]AA[GA]TGATGCGGTTT incorporates four potential nucleotides at position 4 as this accounts for *S. aureus* variants identified via systematic analysis 1,455 complete genomes. For *attL* the patterns include conserved flanking motifs of 5 and 6 bp (CATCA..GATAAG), these motifs are bridged by variable-length sequences ranging from 5 to 15 bp.

Once the *att* sites have been detected further checks are undertaken to confirm that *attR* is identified at the expected location e.g. within 100 bp of *rlmH.* Two modes are available for *rlmH* detection, GFF3-based (parsing the gene annotations) or BLAST-based (querying a bundled 70-species *rlmH* reference database). The BLAST mode enables FASTA-only input, removing the requirement for genome annotation, with users able to provide their own reference *rlmH* gene.

Results are output as a per-genome TSV file recording all detected *att* sites with their positions, patterns and strand orientation, enabling further exploration of the *att* site patterns and genomic location.

**SCC Element Extraction -** sccmec-extract

The extraction module receives *att* site locations and applies a series of biological validation steps to determine whether a complete SCC element can be extracted (Figure 2).

**Figure 2:**
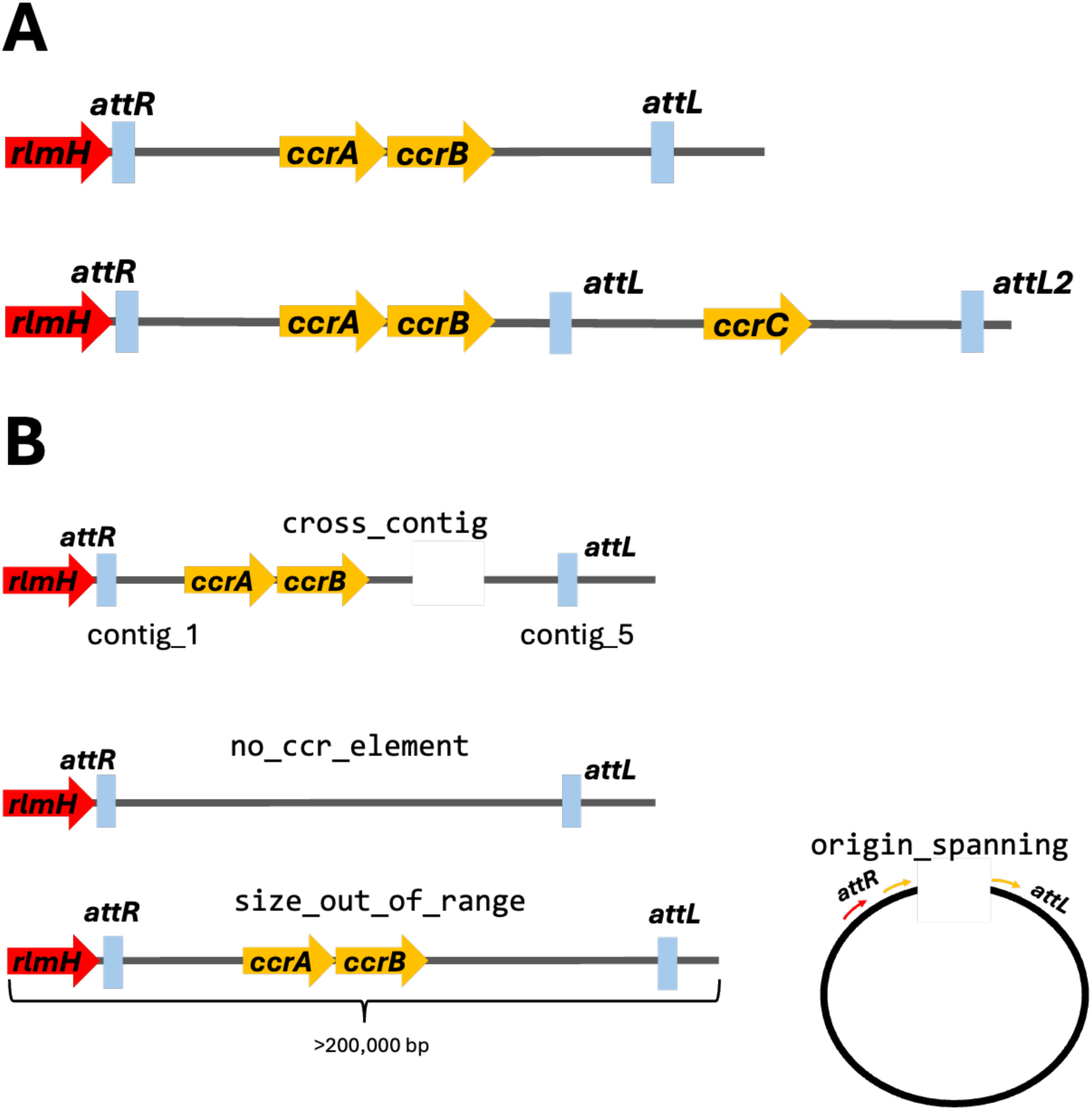
Overview extraction requirements. After obtaining valid *att* sites from `sccmec-locate-att`, `sccmec-extract` performs additional checks to determine if a SCC element can be extracted. **A**. Successful SCC element extraction with *att* sites located on the same contig, *ccr* genes present between *att* sites and *attR* located within proximity of *rlmH*. Note composite elements are also extracted using the *attL* furthest from *attR* as the outer limit. **B.** SCC element failure scenarios: *attR* and *attL* are located on different contigs, `cross_contig`, missing *ccr* genes between *attR/rlmH* and *attL,* `no_ccr_element`, element size is above threshold e.g. 200,000 bp for single element, 250,000 bp for composite elements, `size_out_of_range`, there is also a cross origin check for *att* sites that span the chromosome boundary, `origin_spanning`.

### Same-contig requirement

Both *attR* and *attL* must reside on the same contig. For draft assemblies where SCC elements may span contig boundaries, the tool reports the failure reason (`cross_contig`) rather than attempting contig concatenation, prioritising extraction confidence over completeness.

### *ccr* gene check

After identifying a valid *att* site pair, a BLAST search confirms the presence of *ccr* genes between the *att* sites. Elements with *att* site pairs but no *ccr* genes are classified as `no_ccr_element` (potentially staphylococcal cassette islands, SCI) and recorded in the extraction report but not extracted as FASTA.

### Size filter

Extracted elements must fall between 1,000 bp and 200,000 bp, for single elements, with composite size increased to 250,000 bp for the upper limit. This rejects pattern-overlap artefacts and genome-spanning spurious matches.

### Composite element detection

When multiple *att* site pairs are detected on the same contig, the tool identifies nested or tandem SCC structures. With the `--composite` flag, extraction extends to the outermost *att* site boundary and reports both inner and outer element coordinates, with *ccr* gene checks required within the outer limit to be considered genuine.

### Fallback extraction

When only *attL* is detected (no *attR*), a fallback pathway infers the right boundary from the *rlmH* gene position. This requires: (i) *rlmH* on the same contig as *attL*; (ii) *attL* within 120 kbp of *rlmH*; and (iii) *ccr* genes between *attL* and *rlmH*. This is also reported as `fallback_extracted` alerting users that this path was taken for extraction.

### Origin-spanning detection

Circular chromosomes linearised at the SCC locus produce *att* sites at opposite ends of the sequence, yielding apparent element sizes covering most of the chromosome. When the calculated element size exceeds the maximum but the wrap-around distance (contig length minus element size) falls within a plausible range, the genome is flagged as `origin_spanning` with coordinates for manual extraction.

### Extraction diagnostics

Every genome receives an extraction status and, for failures, a categorised reason: `right_only`, `left_only`, `cross_contig`, `no_sites`, `size_out_of_range`, `no_ccr_element`, or `origin_spanning`. This transparent reporting enables users to understand why extraction failed and to assess whether better assembly quality might resolve the failure or if these are non-addressable failures.

Output includes extracted element FASTA sequences and an 18-column TSV extraction report with comprehensive metadata including element coordinates, size, composite status, contig edge proximity and diagnostic notes. In addition, a separate `ambiguous_att_sites.tsv` file collates all *ccr*-positive non-extracted genomes with their failure reason, detected *att* site positions, *rlmH* contig, element size and explanatory notes. It also includes information on genomes with *attR* and *attL* detected, but no *ccr* genes, as these could contain novel recombinases. This file enables users to investigate extraction failures, for example, identifying cross-contig cases that would be resolved with improved assemblies, examining *att* site pairs that lack *ccr* genes (SCI elements), or locating origin-spanning elements whose coordinates are provided for manual extraction.

**Gene Level Typing -** sccmec-type

Gene-level typing is performed via BLAST against bundled reference databases containing 9 *mec* allotypes (*mecA*, *mecA1*, *mecA2*, *mecB*, *mecC*, *mecC1*, *mecC2*, *mecC3*, *mecD*) and 31 *ccr* allotypes (*ccrA1*-*ccrA14*, *ccrB1*-*ccrB12*, *ccrC1*-*ccrC5*) (Table 1).

**Table 1:** *mec* and *ccr* gene references for gene level typing analysis. SCC*mec*Extractor uses bundled reference databases for *mec* and *ccr* gene detection via BLAST. The table lists all reference allotypes with their GenBank accessions and classification thresholds.

| Reference Gene | Accession | Nucleotide Identity | Nucleotide Coverage |
| --- | --- | --- | --- |
| *mecA* | KC243783.1 | $\geq 95\%$ | $\geq 90\%$ |
| *mecA1* | Y13094.1:2045-4045 | $\geq 95\%$ | $\geq 90\%$ |
| *mecA2* | AM048810.2 | $\geq 95\%$ | $\geq 90\%$ |
| *mecB* | AB498758.1:3738-5762 | ≥95% | ≥90% |
| *mecC* | FR821779.1:35681-37678 | ≥95% | ≥90% |
| *mecC1* | HE993884.1:12033-14029 | ≥95% | ≥90% |
| *mecC2* | KF955540.2:436-2433 | ≥95% | ≥90% |
| *mecC3* | MH155596:119542-121539 | ≥95% | ≥90% |
| *mecD* | KY013611.1:15438-17474 | ≥95% | ≥90% |
| *ccrA1* | AB063171.1:12055-13404 | ≥84.5% | ≥90% |
| *ccrA2* | AB063172.2:9888-11237 | ≥84.5% | ≥90% |
| *ccrA3* | AB014436.1:134-1480 | ≥84.5% | ≥90% |
| *ccrA4* | AF411935.3:7849-9210 | ≥84.5% | ≥90% |
| *ccrA5* | AM904731.1:4627-5976 | ≥84.5% | ≥90% |
| *ccrA6* | MN220716.1:c48983-47634 | ≥84.5% | ≥90% |
| *ccrA7* | AB587080.1:1-1353 | ≥84.5% | ≥90% |
| *ccrA8* | OL880246.1:172-1572 | ≥84.5% | ≥90% |
| *ccrA9* | CP065792.1:c612238-610889 | ≥84.5% | ≥90% |
| *ccrA10* | CP069486.1:c972238-970889 | ≥84.5% | ≥90% |
| *ccrA11* | CP063367.1:c47706-46357 | ≥84.5% | ≥90% |
| *ccrA12* | CP016760.1:c2324024-2322675 | ≥84.5% | ≥90% |
| *ccrA13* | CP065795.1:1089818-1091167 | ≥84.5% | ≥90% |
| *ccrA14* | LT906460.1:c47750-46416 | ≥84.5% | ≥90% |
| *ccrB1* | AB063171.1:13426-15054 | ≥84.5% | ≥90% |
| *ccrB2* | AB063172.2:11259-12887 | ≥84.5% | ≥90% |
| *ccrB3* | AB014436.1:1501-3129 | ≥84.5% | ≥90% |
| *ccrB4* | AF411935.3:9207-10535 | ≥84.5% | ≥90% |
| *ccrB5* | AM904731.1:5997-7625 | ≥84.5% | ≥90% |
| *ccrB6* | AP008934.1:c43044-41416 | ≥84.5% | ≥90% |
| *ccrB7* | AB353724.1:c17096-15468 | ≥84.5% | ≥90% |
| *ccrB8* | AB498756.1:c16657-15029 | ≥84.5% | ≥90% |
| *ccrB9* | OL880246.1:1865-3508 | ≥84.5% | ≥90% |
| *ccrB10* | CP007208.1:2458421-2460049 | ≥84.5% | ≥90% |
| *ccrB11* | LT906460.1:c46380-44755 | ≥84.5% | ≥90% |
| *ccrB12* | CP063367.1:c46337-44709 | ≥84.5% | ≥90% |
| *ccrC1* | AP006716.1:64143-65819 | ≥84.5% | ≥90% |
| *ccrC2* | KR187111.1:30895-32580 | ≥84.5% | ≥90% |
| *ccrC3* | CP065792.1:565984-567660 | ≥84.5% | ≥90% |
| *ccrC4* | CP059679.1:c1833169-1831493 | ≥84.5% | ≥90% |
| *ccrC5* | GU370073.2:c9998-8322 | ≥84.5% | ≥90% |

Novel *mec*: 75-94.9% identity; novel *ccr*: 70-84.4% identity.

Detections are classified into four confidence tiers. For *mec* genes: *full* (>=95% identity, >=90% coverage), *partial* (>=95% identity, 75-90% coverage), *novel_full* (75-94.9% identity, >=90% coverage), and *novel_partial* (75-94.9% identity, 75-90% coverage). For *ccr* genes: *full* (>=84.5% identity, >=90% coverage) and *novel* (70-84.4% identity). Overlapping BLAST hits are resolved using a 500 bp threshold and best bitscore selection.

Detected *ccr* allotype combinations are mapped to IWG-SCC *ccr* complex types 1-22 using a lookup table ^11^. Gene-level typing was chosen over formal SCC*mec* type assignment (types I-XV) because formal types are generally defined for *S. aureus* SCC*mec* elements, rather than SCC*mec* elements carried by non-*aureus* or Mammaliicocci species. Gene-level reporting of *mec* allotypes, *ccr* allotypes and *ccr* complex types is more broadly informative for cross-species work and enables characterisation of novel or divergent elements. Typing can be applied to extracted SCC elements or to whole-genome sequences, enabling gene characterisation regardless of extraction outcome.

**Unified Reporting -** sccmec-report

The `sccmec-report` module merges extraction metadata with typing results into a single unified report TSV. A priority system prefers gene level typing results obtained from extracted SCC elements, with whole-genome typing as a fallback for genomes where extraction failed.

The typing_source column indicates whether typing was performed on the extracted element (sccmec) or the whole genome sequence (wgs).

**Master Pipeline -** sccmec-pipeline

For convenience, `sccmec-pipeline` orchestrates the complete workflow (`locate-att`, `extract`, `type`, `report`) in a single command. It supports batch processing of multiple genomes and both GFF3+FASTA and FASTA-only input modes, with multi-threading.

### Implementation Details

SCC*mec*Extractor is implemented in Python 3.9+ with BioPython and BLAST+ as the only external dependencies. It is available via PyPI (`pip install sccmecextractor`), Docker Hub and Singularity. The source code is hosted on GitHub under an MIT licence.

## Results

### Typing concordance on known SCC*mec* References

SCC*mec*Extractor was evaluated on 32 reference sequences from the `sccmec` tool (v1.2.0) test suite, covering SCC*mec* types I-XV including subtypes (Table 2). Listed expected genes are those used to define the SCC*mec* element type, with SCC*mec*Extractor detecting additional genes when composite elements are present. All 32 references had *ccr* genes correctly detected (100%) as well as *ccr* complex type assignment. Of the 28 complete reference sequences, *mec* genes were detected in 28/28 (100%). Four references (types IIb, IId, IIe, and IVd) are partial sequences deposited in GenBank that do not contain the *mec* complex region; the tool correctly reported no *mec* for these entries while still detecting their *ccr* genes.

**Table 2:**
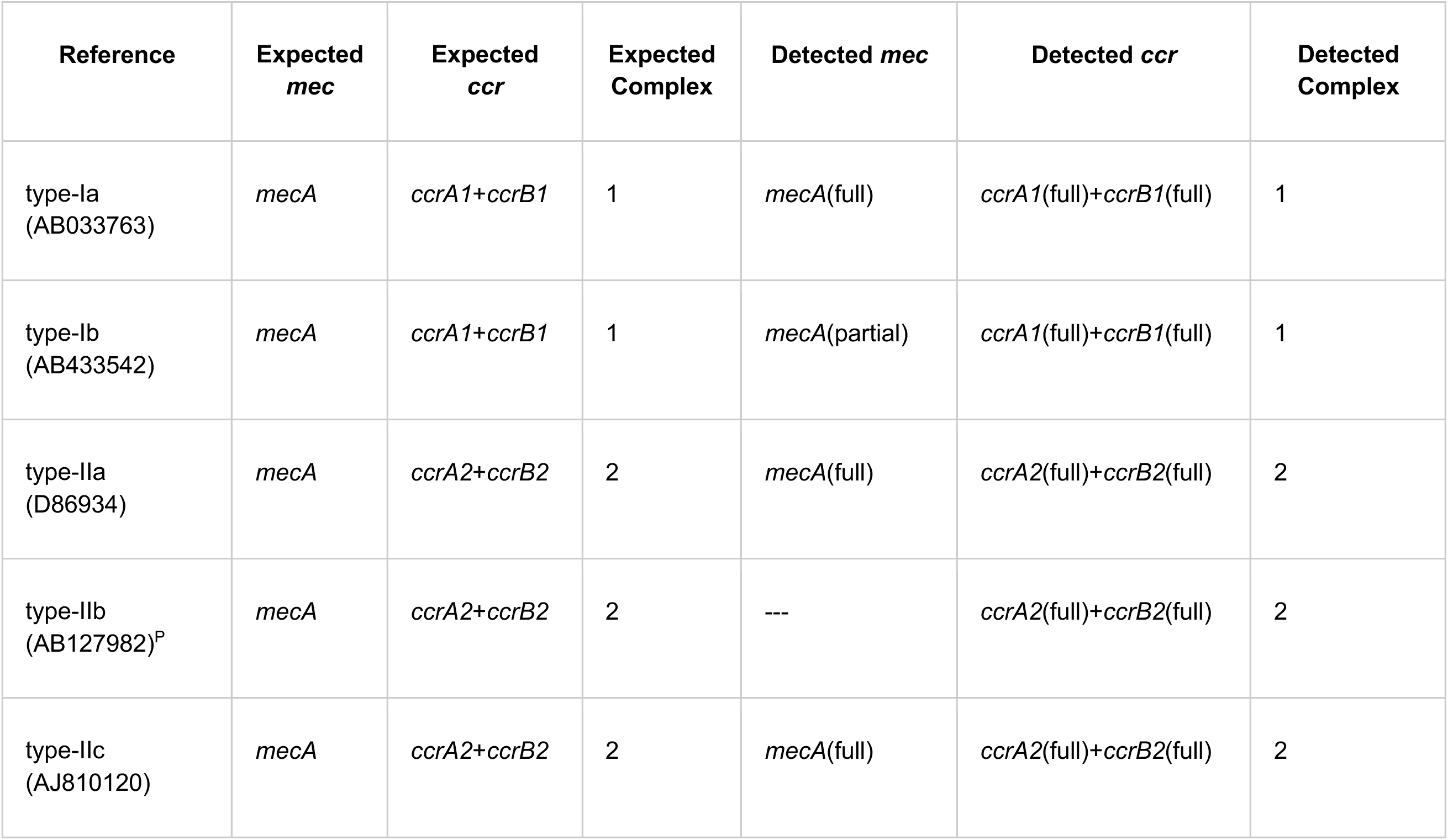
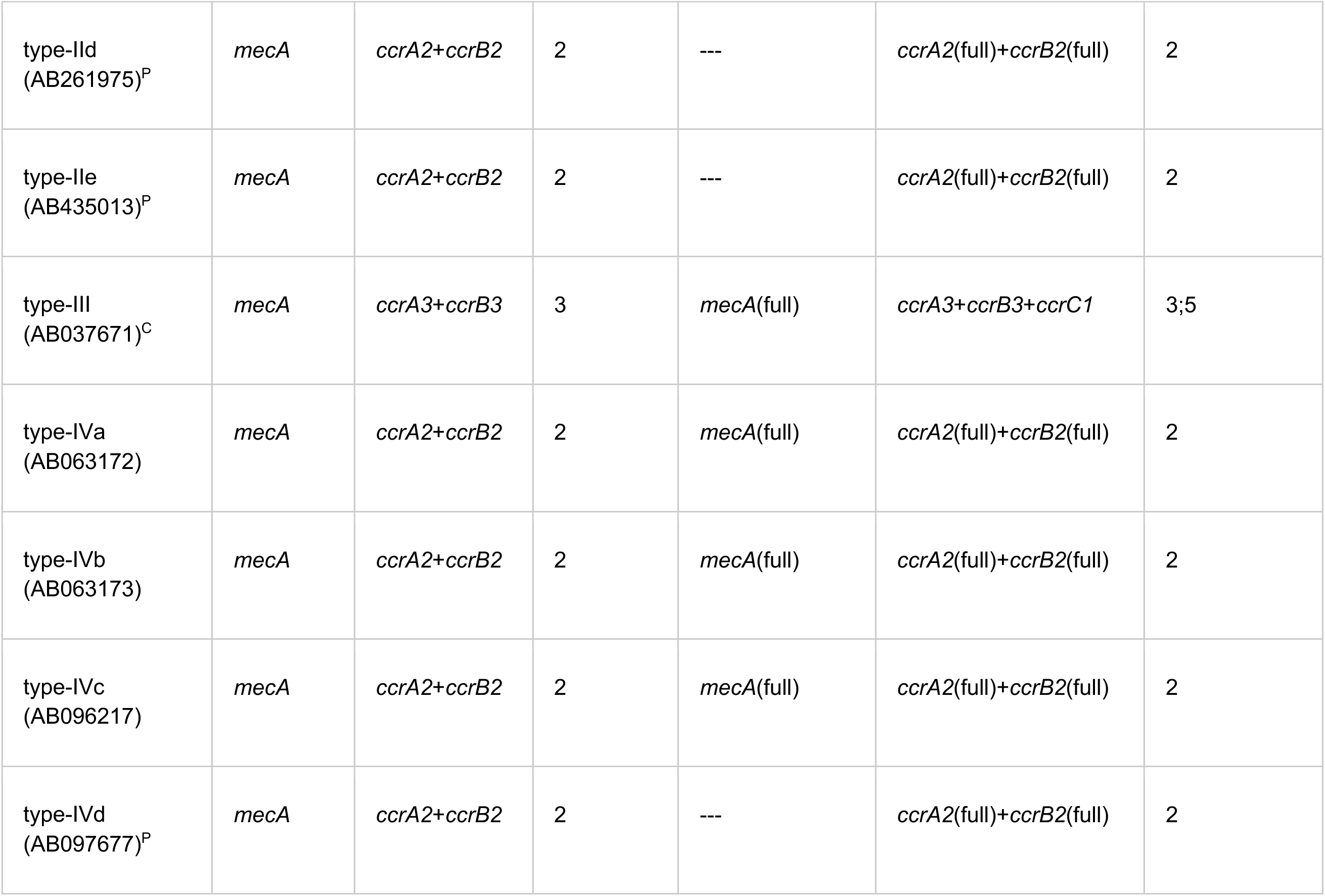
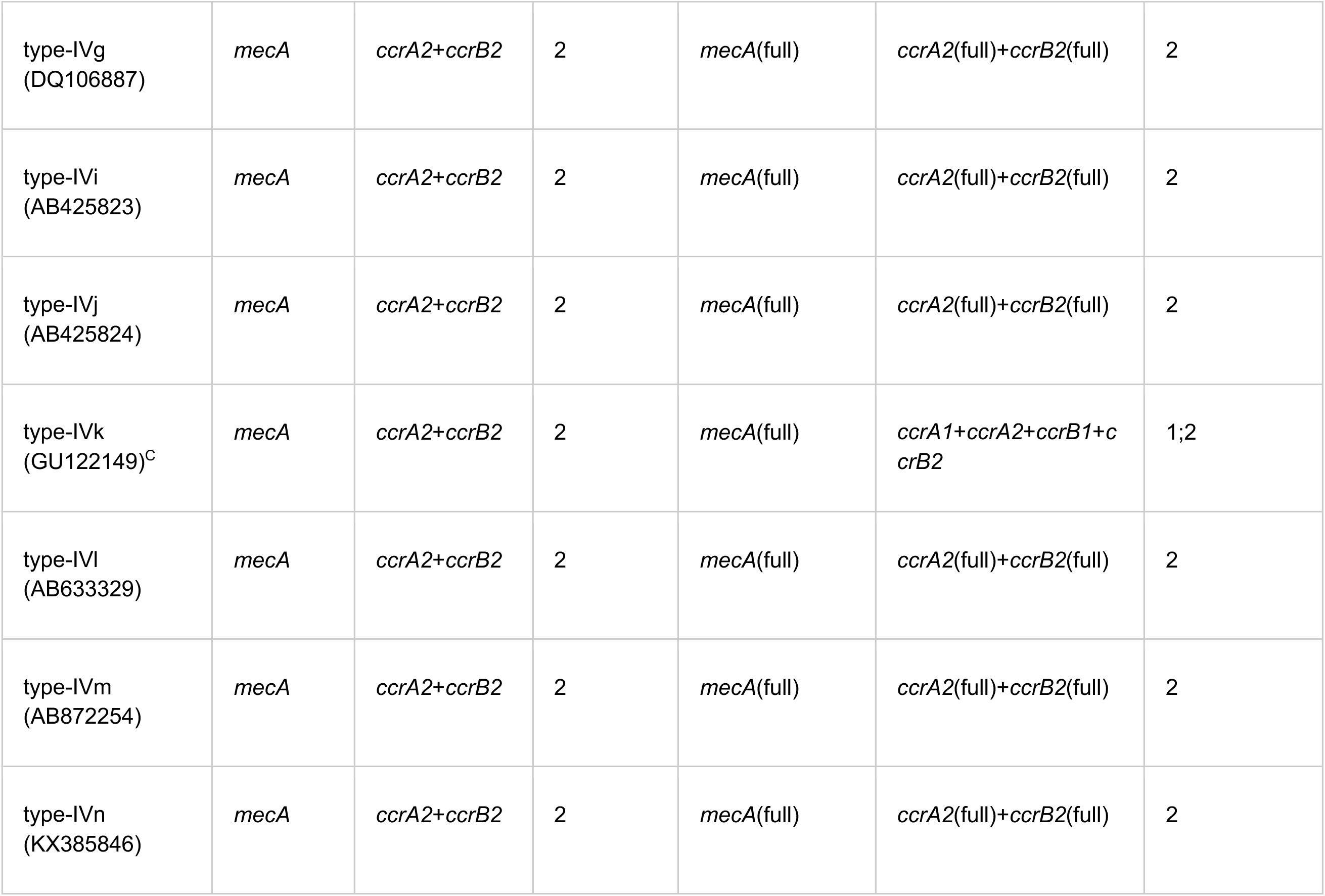
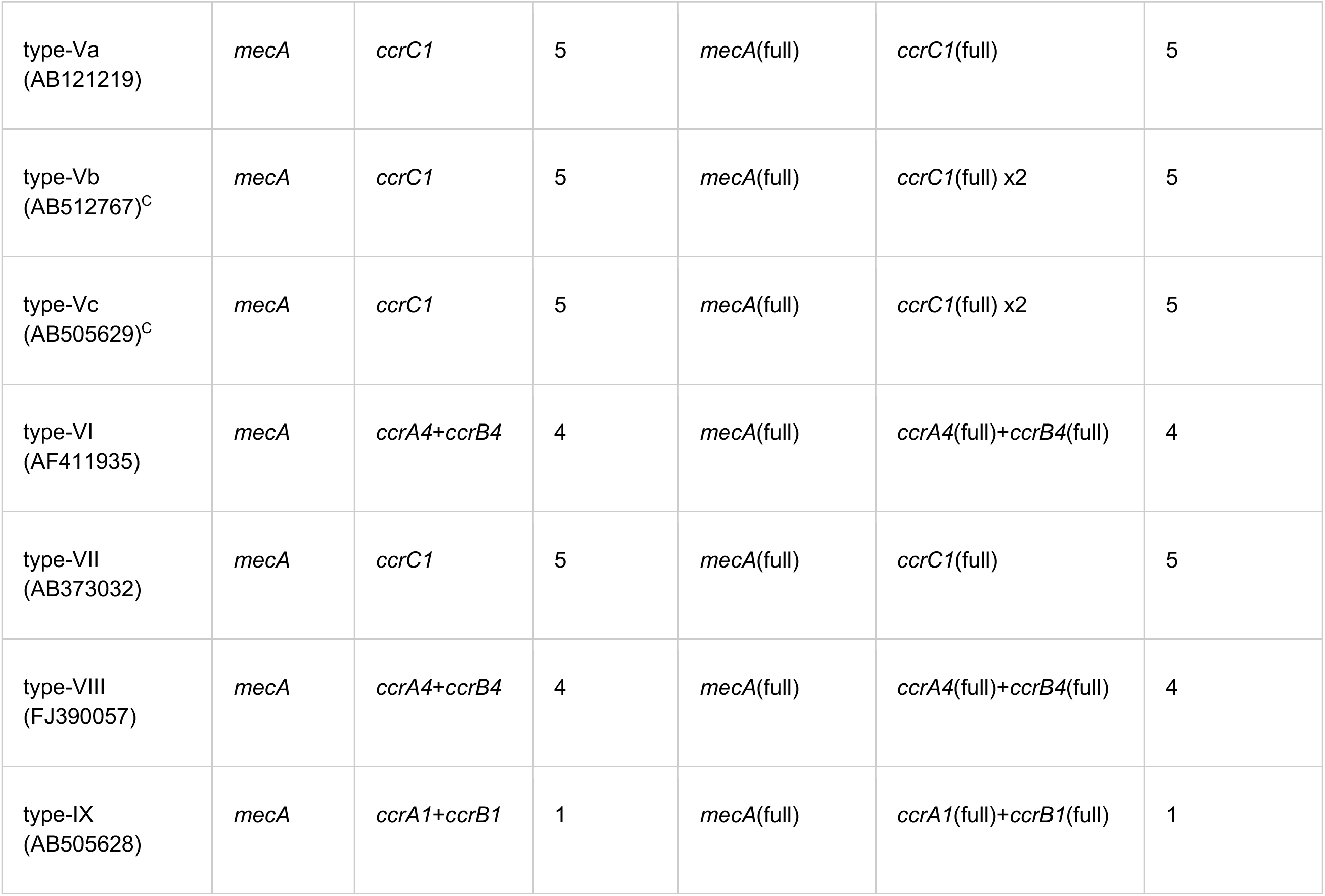
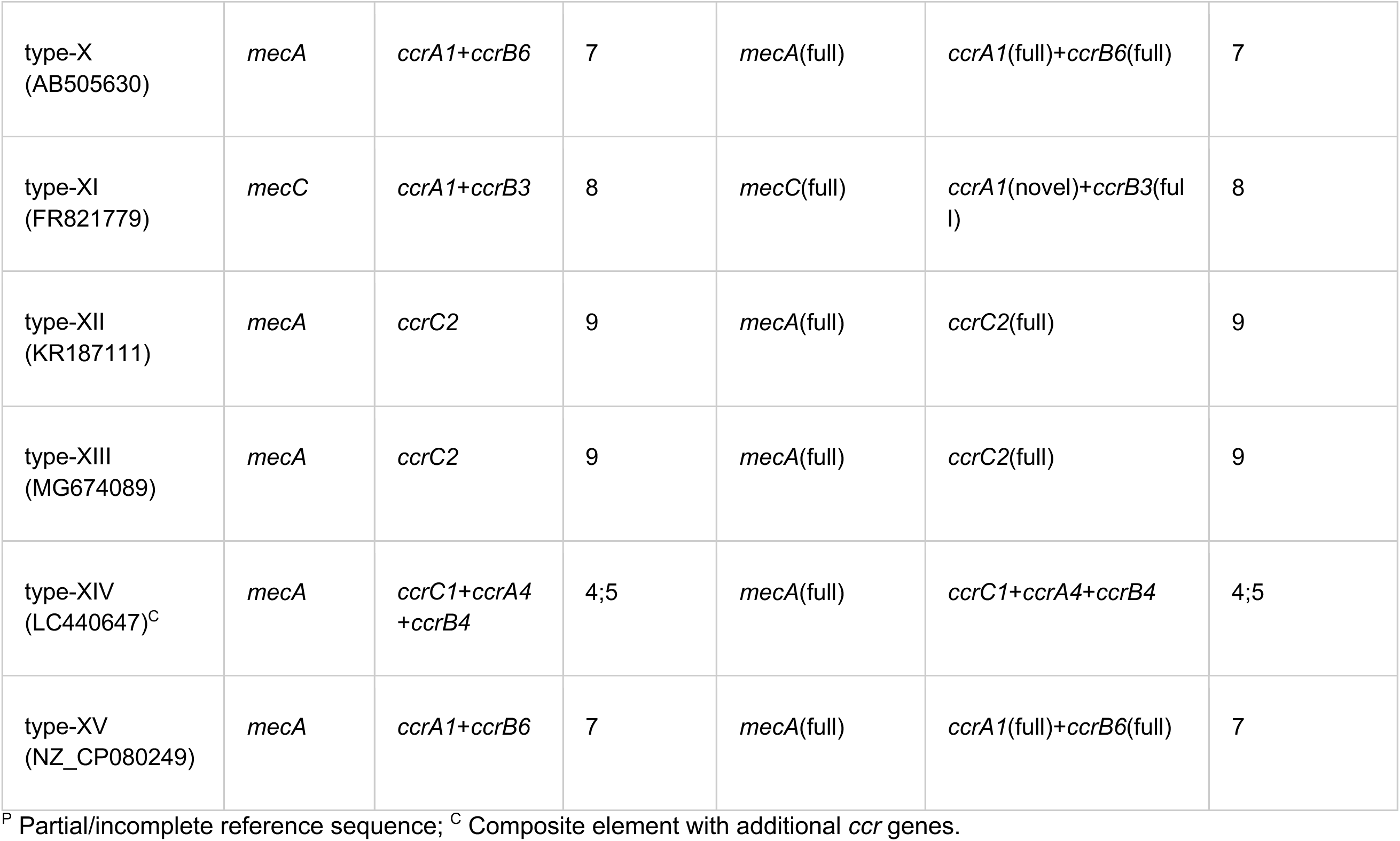
SCC*mec* Type references and typing concordance by SCC*mec*Extractor SCC*mec*Extractor gene detection results on 32 reference sequences from the `sccmec` test suite. Four partial references (marked ^P^) correctly show no *mec* detected because their sequences do not include the *mec* complex region. Composite elements (marked ^C^) show additional *ccr* genes beyond the primary type definition.

SCC*mec*Extractor additionally identified composite elements within reference sequences: type III (AB037671) contained an additional *ccrC1* gene beyond the expected *ccrA3*/*ccrB3*, consistent with the known SCC*Hg* composite structure; type IVk (GU122149) carried additional *ccrA1*/*ccrB1*, indicating a composite element; and type XIV (LC440647) contained *ccrC1* alongside *ccrA4*/*ccrB4*, consistent with its described composite architecture.

### Typing concordance on *S. aureus* whole-genome sequences

To validate typing accuracy at scale, SCC*mec*Extractor was compared against `sccmec` on 1454 *S. aureus* complete genomes from NCBI (Supplementary Table S1). *S. aureus* genomes were obtained by using `datasets`^21^ with `--assembly-level` “complete” or “chromosome”, with the taxon flag `“Staphylococcus aureus”` to ensure contiguous sequence data suitable for SCC*mec* boundary resolution. No filtering was applied based on resistance phenotype, isolation source, or any other biological criterion, though genome GCF_020167475.1 was excluded, as it was misclassified as complete though it contains 102 contigs. However, complete genome assemblies in public databases are enriched for clinically significant strains, which may inflate the proportion of SCC*mec*-carrying genomes relative to the species as a whole. Within this dataset, both tools detected *mecA*/*mecC* in the same 653 genomes (100% concordance) and agreed on *ccr* complex type assignment for all 606 genomes where `sccmec` assigned a single type (100% concordance). An additional 21 genomes assigned “multiple” by `sccmec` also matched SCC*mec*Extractor at the gene level. Runtime was approximately 52 min for SCC*mec*Extractor (∼2.1 s/genome) versus approximately 63 min for `sccmec` (∼2.6 s/genome).

The two tools take complementary approaches that diverge for cases beyond standard *S. aureus* SCC*mec* types. `sccmec` assigns formal SCC*mec* types (I-XV) by matching genome sequences against whole-cassette reference regions, an approach well-suited to epidemiological typing of *S. aureus*. SCC*mec*Extractor takes a gene-level approach, reporting individual *mec* and *ccr* allotypes and their *ccr* complex type, providing a consistent framework that extends to novel gene combinations and non-*aureus* species.

These design differences are apparent in three scenarios. First, SCC*mec*Extractor detected *ccr* genes in 18 genomes carrying novel allotypes (*ccrA5*, *ccrA6*, *ccrA9*, *ccrA11*, *ccrB5*, *ccrB12*, *ccrC4*) absent from `sccmec`‘s targets reference database. Second, SCC*mec*Extractor identified 89 genomes (12.3% of *ccr*-positive) carrying non-*mec* SCC elements, of which `sccmec` correctly detected their *ccr* genes in its targets output but could not assign a type without a *mec* gene, reflecting its design focus on SCC*mec* typing rather than broader SCC diversity. Third, among *mecC*-carrying genomes, SCC*mec*Extractor distinguished specific allotypes (*mecC*, *mecC1*), whereas `sccmec` detected all as “*mecA*+”, sufficient for flagging the presence of a *mec* gene but not for distinguishing *mec* gene variants. This is relevant given the reduced affinity of *mecC* for almost all β-lactam antimicrobials^22^.

The approaches also differ in their handling of composite (tandem/nested) elements. SCC*mec*Extractor identifies potential composite SCC elements via gene-level typing, with 72 *S. aureus* genomes carrying multiple *ccr* complex types. SCC*mec*Extractor then confirms composite structure before extraction, where multiple *attL* sites are flagged as inner, closest to *attR* and outer, further from *attR*. A composite element is then defined when there are *ccr* complexes between both the inner *attL* and *attR*, and the inner and outer *attL*, as shown in Figure 2. This resulted in 5 composite SCC element extractions out of 72 genomes carrying multiple *ccr* complex types, representing 0.8% of the total extracted SCC elements for *S. aureus*. Whereas `sccmec` flagged 105 genomes with a “multiple” subtype, indicating matches to more than one reference cassette. Exploring this further reveals these represent three distinct situations: 67 (64%) were subtype ambiguity within a single type (e.g. simultaneous IVa and IVn matches to a single element), 18 (17%) reflected partial sequence similarity to a second reference cassette despite carrying only one *ccr* complex and 20 (19%) carried genuinely distinct *ccr* complex types. With SCC*mec*Extractor’s gene-level typing, 72 multi-*ccr* complex genomes were identified compared to `sccmec`’s 20 genuine multi-type calls. Gene synteny analysis provides further validation for extracted elements and distinguishes between tandem integration events from single integration events for SCC elements that carry multi-*ccr* complexes. Overall, these results illustrate how whole-cassette reference matching and *att* site-based boundary detection address complementary aspects of SCC biology.

### SCC element extraction across staphylococci and *Mammaliicoccus*

The full SCC*mec*Extractor pipeline was applied to three datasets (Supplementary Table S1), all using BLAST-based *rlmH* detection in FASTA-only mode and FASTA + GFF3 mode, though for benchmarking the BLAST-based results were used. The datasets included 1,454 *S. aureus* complete genomes, 5,295 non-*aureus Staphylococcus* genomes from 63 species and 548 *Mammaliicoccus* genomes from 6 species. Extraction performance is summarised in Table 3. As with the *S. aureus* dataset, non-*aureus* staphylococci and *Mammaliicoccus* genomes were obtained using `datasets`.

**Table 3:** SCC element extraction performance across three datasets. Summary of SCC element extraction using the full SCC*mec*Extractor pipeline across *S. aureus*, non-*aureus Staphylococcus* and *Mammaliicoccus*.

| Metric | *S. aureus* | Non-*aureus Staphylococcus* | *Mammaliicoccus* | Combined |
| --- | --- | --- | --- | --- |
| Total genomes | 1,454 | 5,295 | 548 | 7,297 |
| Species | 1 | 63 | 6 | 70 |
| *ccr*-positive genomes | 726 (49.9%) | 2,918 (55.1%) | 269 (49.1%) | 3,913 (53.6%) |
| **Total extracted** | **630 (43.3%)** | **828 (15.6%)** | **104 (19.0%)** | **1,562 (21.4%)** |
| *ccr*+ extraction rate | 86.8% | 28.3% | 38.6% | 39.9% |
| Effective extraction rate | 87.3% | 58.8% | 61.9% | - |
| **Element classification** |  |  |  |  |
| SCC<sub>mec</sub> (*mec* + *ccr*) | 553 (87.8%) | 368 (44.4%) | 25 (24.0%) | 946 (60.6%) |
| Non-*mec* SCC (*ccr*, no *mec*) | 77 (12.2%) | 460 (55.6%) | 79 (76.0%) | 616 (39.4%) |
| **Features** |  |  |  |  |
| Composite extractions | 5 (0.8%) | 76 (9.2%) | 11 (10.6%) | 92 (5.9%) |
| Fallback extractions | 11 (1.7%) | 37 (4.5%) | 12 (11.5%) | 60 (3.8%) |
| No-*ccr* elements (SCI) | 94 | 284 | 72 | 450 |
| Origin-spanning detected | 2 | 3 | 0 | 5 |
| **Failure analysis (*ccr*+)** |  |  |  |  |
| Assembly-limited failures | 2 | 1,452 (69.5%) | 100 (60.6%) | 1554 |
| Right_only (no *attL*) | 66 | 477 | 49 | 592 |
| Addressable failures | 80 | 545 | 60 | 685 |
| Median (bp) | 29,415 | 31,557 | 37,326 | - |
| Range (bp) | 12,109-99,888 | 11,153-158,206 | 14,630-107,704 | - |

Effective extraction rate excludes assembly-limited genomes (cross-contig, origin-spanning, *attL/rlmH* on different contigs) and *ccr* genes without a valid pair e.g. lone *ccrA* without a corresponding *ccrB*. *right\_only* counts include *attB* overlap artefacts (31 non- *aureus*, 9 *Mammaliicoccus*) where the *attL* regex matched the empty chromosomal integration site rather than a genuine element boundary.

Across all three datasets, 1,562 SCC elements were extracted from 7,297 genomes spanning 70 species and two genera (Supplementary Table S3). The raw extraction rate among *ccr*-positive genomes was 86.8% for *S. aureus*, 28.3% for non-*aureus Staphylococcus* and 38.6% for *Mammaliicoccus*. The lower non-*aureus* and *Mammaliicoccus* rates reflect the predominance of draft assemblies. To assess tool performance independent of assembly quality, we calculated an effective extraction rate by excluding genomes where extraction is impossible regardless of tool capability: assembly-limited genomes (those with *att* sites on different contigs, origin-spanning, or *attL*/*rlmH* on different contigs) and genomes with unpaired *ccr* genes (lone *ccrA* or *ccrB* without a functional recombinase pair). The effective extraction rate was 87.3% (630/722) for *S. aureus*, 58.8% (828/1,408) for non-*aureus Staphylococcus*, and 61.9% (104/168) for *Mammaliicoccus*. Filtering the extraction results based on species revealed some species within the non-*aureus* dataset had effective extraction rates of 100.0%, with clinically relevant species such as *S. saprophyticus, S. lugdunensis,* and *S. pseudintermedius* had rates of 94.9%, 92.9% and 70.9%, respectively (Supplementary Table S4) Whereas some clinically relevant species had lower effective extraction rates than the average e.g. *S. epidermidis* 57.5%, or slightly higher as in the case of *S. haemolyticus* at 62.9%.

#### Failure analysis

Assembly fragmentation was the dominant cause of extraction failure across all datasets. For *S. aureus* (all complete genomes), cross-contig failures were absent; the 96 *ccr*-positive non-extracted genomes comprised of 66 `right_only` (no matching *attL* pattern), 12 classified as `no_ccr_element` (*ccr* detected outside element bounds), 9 `left_only` (no *ccr* genes found between *attL* and *rlmH*), 4 with no *att* sites detected, 2 `origin_spanning` genomes (flagged with wrap-around coordinates), 2 with unpaired *ccr* genes (lone *ccrA* without *ccrB* partner) and 1 element `size_out_of_range`. For non-*aureus Staphylococcus*, 1,452 of 2,090 *ccr*-positive failures (69.5%) were assembly-limited: 1,270 `cross-contig`, 175 with *attL* and *rlmH* on different contigs, 4 with no *rlmH* detected (due to the gene being split over two contigs) and 3 `origin-spanning`. The remaining 545 addressable failures included 477 genomes requiring novel *attL* patterns e.g. `right_only` failures, where extreme bridge sequence diversity (45+ unique bridges) yielded diminishing returns for further pattern expansion. Another 33 with `size_out_of_range`, 25 with no *att* sites detected and 10 `left_only`. For *Mammaliicoccus*, 100 of 165 *ccr*-positive failures (60.6%) were assembly-limited (93 `cross-contig`, 6 with *attL* and *rlmH* on different contigs, 1 with no *rlmH* detected as the gene was split between two contigs), leaving 60 addressable failures including 49 `right_only`.

#### Composite elements

Composite elements were detected in 5 (0.8%) of *S. aureus* extractions, 76 (9.2%) of non-*aureus* and 11 (10.6%) of *Mammaliicoccus* extractions.

#### Fallback extraction

SCC elements were recovered for 11 *S. aureus* (1.7% of extractions), 37 non-*aureus* (4.5%) and 12 *Mammaliicoccus* (11.5%) genomes. The increasing fallback rate across datasets reflects growing *attR* site divergence away from the *Staphylococcus*-derived patterns.

#### No-*ccr* elements

The *ccr* gene check correctly classified 450 *att* site pairs across all datasets as `no_ccr_element` (SCI): 94 in *S. aureus*, 284 in non-*aureus Staphylococcus* and 72 in *Mammaliicoccus*. In *Mammaliicoccus*, 63 of 72 SCI elements (87.5%) carried *mec* genes (predominantly *mecA1*), consistent with the established role of *M. sciuri* group species as the evolutionary reservoir of *mecA*-family genes^23^, where native *mec* homologues are chromosomally encoded independently of SCC*mec*.

### SCC element diversity beyond SCC*mec*

A striking gradient in element composition emerged across the three datasets (Table 3). In *S. aureus*, 87.8% (553/630) of extracted elements were SCC*mec* (carrying both *mec* and *ccr*) and only 12.2% (77/630) were non-*mec* SCC. In non-*aureus Staphylococcus*, there were more non-*mec* SCC elements, at 55.6% (460/828) vs 44.4% (368/828). In *Mammaliicoccus*, non-*mec* SCC dominated at 76.0% (79/104), the highest proportion within our datasets.

This gradient reflects both MRSA-centric bias in *S. aureus* research and the genuine prevalence of non-*mec* SCC elements outside *S. aureus*. Among the 2,918 *ccr*-positive non-*aureus* genomes, 1,295 (44.4%) of those genomes carried *ccr* without any detectable *mec* gene. Extracted non-*mec* SCC carriage varied dramatically by species: environmental and commensal species such as *S. equorum*, *S. chromogenes*, *S. xylosus* at 100% carried predominantly non-*mec* SCC, while clinical species such as *S. epidermidis* (42.2% non-*mec* SCC) and *S. pseudintermedius* (7.6% non-*mec* SCC) carried mostly SCC*mec* (Supplementary Table S5).

#### Novel *ccr* diversity

In non-*aureus* staphylococci, 120 genomes (4.1%) carried novel *ccr* allotypes below the 84.5% identity threshold for confirmed assignment^11^. In *Mammaliicoccus*, novel *ccr* diversity was far greater: 46.5% of *ccr*-positive genomes carried novel *ccr* complex types, with *ccr* complex types 20 (*ccrA9*/*ccrB5*) and 21 (*ccrC4*) being the most common, types rare or absent in the *S. aureus* dataset.

#### *Mammaliicoccus* gene carriage

The *Mammaliicoccus* gene carriage pattern differed markedly from *Staphylococcus*. *mecA1* (not *mecA*) was the dominant *mec* gene, detected in 321/548 genomes (58.6%), supporting the hypothesis that *M. sciuri* group species represent the evolutionary origin of *mecA*. Strikingly, 42.2% of genomes carried *mec* without detectable *ccr*, suggesting widespread chromosomal *mec* integration outside SCC context.

### Comparison with existing tools

Table 4 summarises the feature comparison between SCC*mec*Extractor and existing tools. Typing concordance with `sccmec` was demonstrated on both reference sequences (32/32 correct *ccr* complex types) and *S. aureus* whole genomes (100% concordance on 1,454 genomes).

**Table 4.**
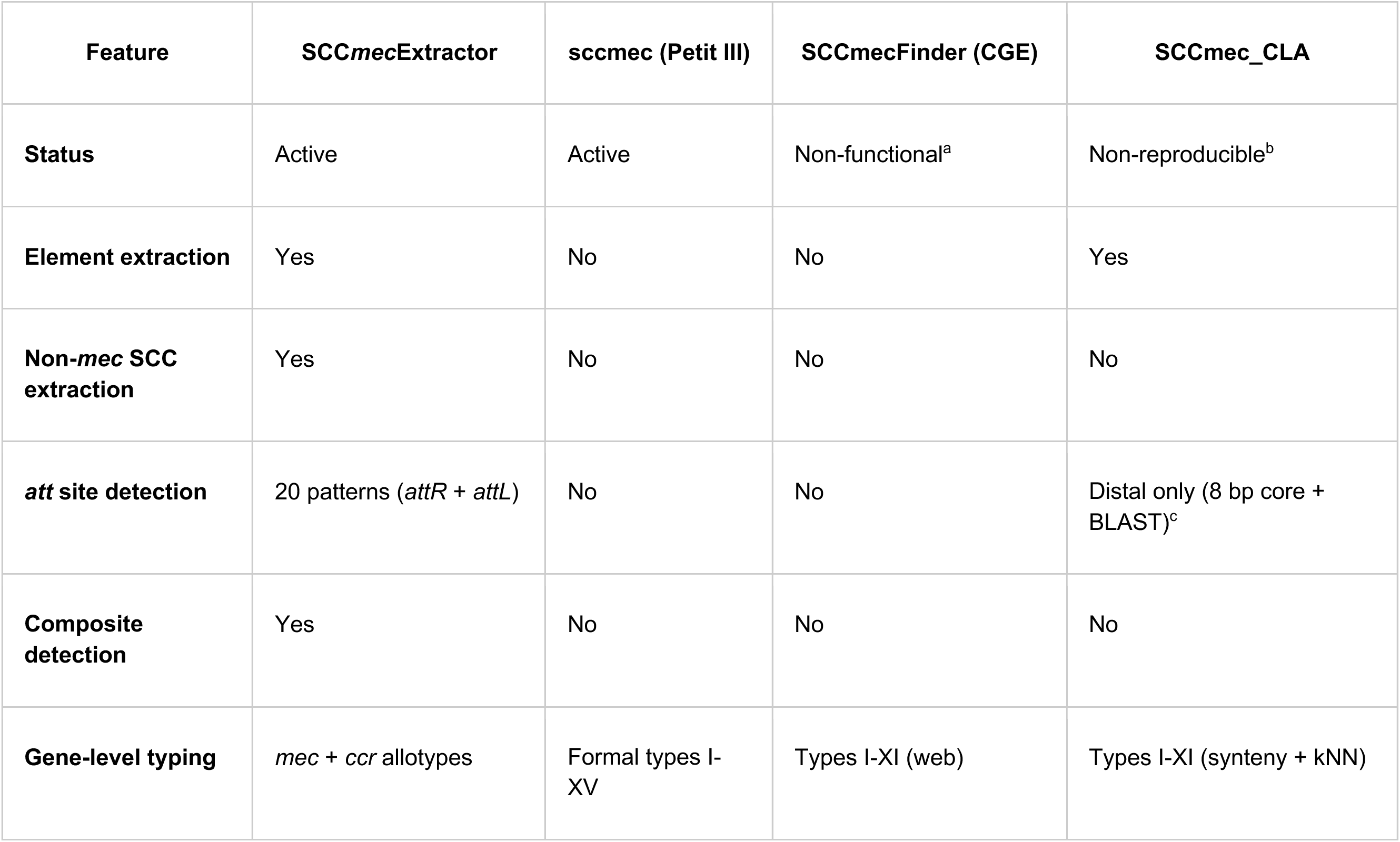
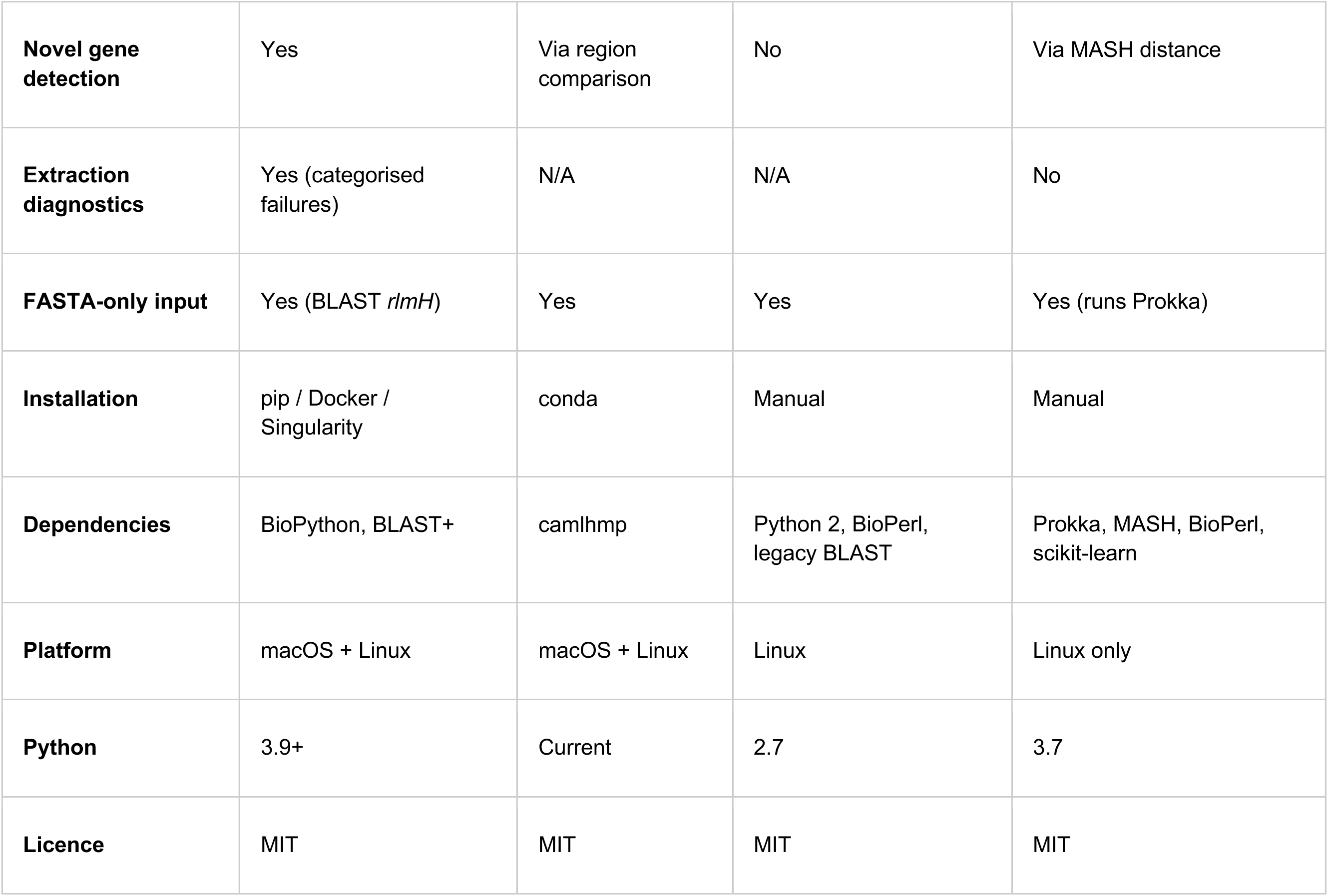
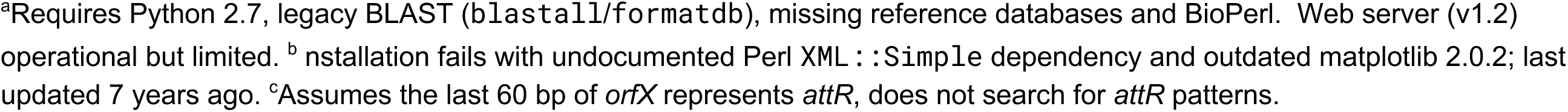
Feature comparison with existing SCC*mec* tools.

| Feature | SCCmecExtractor | sccmec (Petit III) | SCCmecFinder (CGE) | SCCmec_CLA |
| --- | --- | --- | --- | --- |
| **Status** | Active | Active | Non-functional<sup>a</sup> | Non-reproducible<sup>b</sup> |
| **Element extraction** | Yes | No | No | Yes |
| **Non-mec SCC extraction** | Yes | No | No | No |
| **att site detection** | 20 patterns (*attR* + *attL*) | No | No | Distal only (8 bp core + BLAST)<sup>c</sup> |
| **Composite detection** | Yes | No | No | No |
| **Gene-level typing** | *mec* + *ccr* allotypes | Formal types I-XV | Types I-XI (web) | Types I-XI (synteny + kNN) |
| **Novel gene detection** | Yes | Via region comparison | No | Via MASH distance |
| **Extraction diagnostics** | Yes (categorised failures) | N/A | N/A | No |
| **FASTA-only input** | Yes (BLAST *rlmH*) | Yes | Yes | Yes (runs Prokka) |
| **Installation** | pip / Docker / Singularity | conda | Manual | Manual |
| **Dependencies** | BioPython, BLAST+ | camlhmp | Python 2, BioPerl, legacy BLAST | Prokka, MASH, BioPerl, scikit-learn |
| **Platform** | macOS + Linux | macOS + Linux | Linux | Linux only |
| **Python** | 3.9+ | Current | 2.7 | 3.7 |
| **Licence** | MIT | MIT | MIT | MIT |

<sup>a</sup>Requires Python 2.7, legacy BLAST (blastall/formatdb), missing reference databases and BioPerl. Web server (v1.2) operational but limited. <sup>b</sup> nstallation fails with undocumented Perl XML : : Simple dependency and outdated matplotlib 2.0.2; last updated 7 years ago. <sup>c</sup>Assumes the last 60 bp of *orfX* represents *attR*, does not search for *attR* patterns.

SCC*mec*Finder (CGE) could not be evaluated as a standalone tool. The software requires Python 2.7, legacy BLAST (`blastall`/`formatdb`), gene databases not included in the download and BioPerl; it is effectively non-functional for local benchmarking. The CGE web server (v1.2) remains operational but is limited to SCC*mec* types I-XI and does not perform extraction.

SCCmec_CLA could not be evaluated as installation failed on a standard Linux HPC following the repository’s instructions, with two independent errors: an undocumented Perl `XML::Simple` dependency required by the bundled PROKKA 1.12, as well as an outdated matplotlib 2.0.2 requirement that fails to compile without system-level library access. The tool was last updated seven years ago, uses Python 3.7 and bundles a Linux-only Mash binary. However, reviewing the code confirmed that it locates the distal *att* (referred to here as *attL* but by SCCmec_CLA as *attR*) site boundary using the 8 bp core motif of the *att* sequences, with BLAST validation against a reference database of known *att* sites. The approximation of the *attR* (referred to as *attL* by SCCmec_CLA) within *rlmH* is reasonable, though this results in no specific *attR* sequence being reported. Finally, the element typing is a thorough approach, examining synteny of the *mec* complex, identifying *ccr* allotypes and using k-nearest-neighbour subtype prediction against reference cassettes, however typing only occurs on extracted sequences, therefore if extraction fails no whole genome typing occurs.

## Discussion

SCC*mec*Extractor fills a bioinformatic gap for staphylococcal genomics by combining SCC element extraction, gene-level typing and non-*mec* SCC detection in a single package. Benchmarking across 7,297 genomes from 70 species and two genera demonstrates that it performs accurately and reveals biological diversity invisible to existing tools.

### Non-*mec* SCC detection is a unique and biologically significant capability

The progressive increase in non-*mec* SCC prevalence from *S. aureus* (12.2%) to non-*aureus Staphylococcus* (55.6%) to *Mammaliicoccus* (76.0%) reveals that non-*mec* SCC elements are the dominant SCC class outside *S. aureus*. These elements carry *ccr* recombinases and integrate at *rlmH* but lack methicillin resistance genes, with their function remaining largely uncharacterised. However, their prevalence, particularly in environmental and commensal species, suggests they play important roles in staphylococcal adaptation beyond antibiotic resistance. The extraction of 616 non-*mec* SCC sequences across this study provides material for downstream functional analysis.

### Gene-level typing is taxonomically appropriate for cross-species work

Formal SCC*mec* types I-XV are defined by IWG-SCC specifically for *S. aureus*. Applying these types to non-*aureus* species is neither appropriate nor informative, as many non-*aureus* species carry *ccr* allotypes and combinations not represented in the *S. aureus* typing scheme. Gene-level reporting of *mec* allotypes, *ccr* allotypes and *ccr* complex types provides a consistent framework for characterising SCC elements regardless of host species, while remaining compatible with formal typing for *S. aureus* isolates.

### Cross-genus applicability extends the tool beyond *Staphylococcus*

The successful extraction of 104 SCC elements from *Mammaliicoccus* (formerly the *S. sciuri* group^24^) demonstrates that SCC*mec*Extractor can be applied to related genera with the addition of genus-specific *rlmH* sequences to the reference database. The *Mammaliicoccus* analysis revealed that *mecA1*, not *mecA* is the dominant *mec* gene in this genus and that nearly half of *ccr*-positive genomes carry novel *ccr* complex types not represented in existing typing schemes. These findings support the use of genus-wide SCC analysis for understanding the evolutionary origins and diversity of SCC elements.

### Composite element detection reveals tandem SCC integration across species

Across all datasets, 92 of 1,562 extracted elements (5.9%) were confirmed composite structures containing nested or tandem SCC elements, spanning 20 species across both genera. Composite prevalence increased outside *S. aureus* (0.8%) to non-*aureus* staphylococci (9.2%) and *Mammaliicoccus* (10.6%), mirroring the broader trend of greater SCC diversity in non-*aureus* species. The make-up of these composite elements also followed this pattern: all 5 *S. aureus* composites were SCC*mec*, while 59.2% of non-*aureus* composites (45/76) were non-*mec* SCC. Composite confirmation requires identification of internal *att* site boundaries and verification of *ccr* gene content in the outer region, a capability unique to extraction-based approaches. Reference-matching tools can identify genomes with multiple *ccr* complex types but cannot distinguish genuine tandem element integration from multi-*ccr* single elements or subtype ambiguity.

### Assembly quality is the primary extraction limitation

The same-contig requirement, mandating that both *att* sites reside on a single contig, is a deliberate design choice that prioritises extraction confidence. Assembly fragmentation accounted for 69.5% of *ccr*-positive failures in non-*aureus* staphylococci and 60.6% in *Mammaliicoccus*. This is a field-wide challenge inherent to short-read draft assemblies, not a limitation specific to SCC*mec*Extractor. Cross-referencing *ccr* gene locations with *att* site coordinates revealed that an additional 39 non-aureus genomes classified as `left_only` failures had *ccr* genes co-located with *attL* sites on a different contig from *rlmH*, suggesting the true proportion of assembly-limited failures is higher than the current classification captures. The *ccr* and *mec* gene location columns in the typing output facilitate manual review of such ambiguous cases. Long-read sequencing technologies that produce more contiguous assemblies would substantially improve extraction rates. SCCmec_CLA addresses this by concatenating contigs, but this approach introduces uncertainty about element integrity that we consider unacceptable for confident downstream analysis. Our strategy is to supply the boundary location to users, who can then carry out more specific approaches e.g. PCR and Sanger sequencing to bridge the contig gaps, to resolve whole SCC elements.

### FASTA-only mode removes annotation dependency

The BLAST-based *rlmH* detection mode, validated against a 70-species reference database, enables SCC*mec*Extractor to process raw assemblies without requiring Bakta, Prokka, or other annotation tools. This simplifies integration into routine genomic surveillance pipelines. Furthermore, users can supply their own reference *rlmH* genes for diverse isolates not covered by SCC*mec*Extractor *rlmH* database.

### Limitations

SCC*mec*Extractor has several limitations. First, the same-contig requirement means that SCC elements spanning contig boundaries cannot be extracted; this is the primary constraint for draft assemblies. The `ambiguous_att_sites.tsv` output provides per-genome *att* site coordinates and failure categorisation to facilitate manual investigation of these cases. Second, the tool requires *rlmH* to anchor the *attR* site; the 70-species reference database covers the genus well, but novel or highly divergent species may require a custom reference via the `--rlmh-ref` option. Genome annotation would overcome this limitation. Third, the current 20 *att* site patterns cover the majority of known SCC elements, but extreme *attL* bridge diversity across non-*aureus* species (45+ unique bridges identified) means some species-specific variants remain unrepresented. Fourth, SCC*mec*Extractor does not classify *mec* gene complex class (A-E), which would require detection of regulatory genes (*mecR1*, *mecI*) and insertion sequences (IS431, IS1272) and does not perform J region subtyping.

### Future directions

Planned extensions include IS element detection to enable *mec* complex class assignment, expanded *att* site patterns as more SCC diversity is characterised through long-read sequencing and formal SCC*mec* type annotation as an optional output for isolates.

## Acknowledgements

I’d like to thank Dr Gavin Paterson for his interest in SCC*mec* elements which led me to explore these elements from non-*aureus* sources and encouraged me to create this tool.

## References

1. Petit III, R., A. sccmec.

2. Matuszewska, M., Murray, G. G. R., Harrison, E. M., Holmes, M. A. & Weinert, L. A. The Evolutionary Genomics of Host Specificity in Staphylococcus aureus. Trends in Microbiology 28, 465–477 (2020).

3. Blane, B. et al. Evaluating the impact of genomic epidemiology of methicillin-resistant Staphylococcus aureus (MRSA) on hospital infection prevention and control decisions. Microbial Genomics 10, 001235 (2024).

4. Ubukata, K., Nonoguchi, R., Matsuhashi, M. & Konno, M. Expression and inducibility in Staphylococcus aureus of the mecA gene, which encodes a methicillin-resistant S. aureus-specific penicillin-binding protein. Journal of Bacteriology 171, 2882–2885 (1989).

5. García-Álvarez, L. et al. Meticillin-resistant Staphylococcus aureus with a novel mecA homologue in human and bovine populations in the UK and Denmark: a descriptive study. The Lancet Infectious Diseases 11, 595–603 (2011).

6. Ito, T. et al. Structural Comparison of Three Types of Staphylococcal Cassette Chromosome mec Integrated in the Chromosome in Methicillin-Resistant Staphylococcus aureus. Antimicrobial Agents and Chemotherapy 45, 1323–1336 (2001).

7. Boundy, S. et al. Characterization of the Staphylococcus aureus rRNA Methyltransferase Encoded by orfX, the Gene Containing the Staphylococcal Chromosome Cassette mec (SCCmec) Insertion Site. J Biol Chem 288, 132–140 (2013).

8. Wang, L., Ahmed, M. H., Safo, M. K. & Archer, G. L. A Plasmid-Borne System To Assess the Excision and Integration of Staphylococcal Cassette Chromosome *mec* Mediated by CcrA and CcrB. J Bacteriol 197, 2754–2761 (2015).

9. Misiura, A. et al. Roles of two large serine recombinases in mobilizing the methicillin-resistance cassette SCC *mec*: Roles of the SCCmec recombinases. Molecular Microbiology 88, 1218–1229 (2013).

10. Wang, L., Safo, M. & Archer, G. L. Characterization of DNA Sequences Required for the CcrAB-Mediated Integration of Staphylococcal Cassette Chromosome *mec*, a Staphylococcus aureus Genomic Island. J Bacteriol 194, 486–498 (2012).

11. Huang, J. et al. Unearthing New ccr Genes and Staphylococcal Cassette Chromosome Elements in Staphylococci Through Genome Mining. The Journal of Infectious Diseases jiae044 (2024) doi:10.1093/infdis/jiae044.

12. Lakhundi, S. & Zhang, K. Methicillin-Resistant Staphylococcus aureus: Molecular Characterization, Evolution, and Epidemiology. Clin Microbiol Rev 31, e00020–18 (2018).

13. MacFadyen, A. C. et al. A mecC allotype, mecC3, in the CoNS Staphylococcus caeli, encoded within a variant SCCmecC. Journal of Antimicrobial Chemotherapy 74, 547–552 (2019).

14. International Working Group on the Classification of Staphylococcal Cassette Chromosome Elements (IWG-SCC). Classification of staphylococcal cassette chromosome mec (SCCmec): guidelines for reporting novel SCCmec elements. Antimicrob Agents Chemother 53, 4961–4967 (2009).

15. Urushibara, N., Aung, M. S., Kawaguchiya, M. & Kobayashi, N. Novel staphylococcal cassette chromosome mec (SCCmec) type XIV (5A) and a truncated SCCmec element in SCC composite islands carrying speG in ST5 MRSA in Japan. Journal of Antimicrobial Chemotherapy dkz406 (2019) doi:10.1093/jac/dkz406.

16. Wang, W. et al. Novel SCCmec type XV (7A) and two pseudo-SCCmec variants in foodborne MRSA in China. Journal of Antimicrobial Chemotherapy 77, 903–909 (2022).

17. Luong, T. T., Ouyang, S., Bush, K. & Lee, C. Y. Type 1 Capsule Genes of Staphylococcus aureus Are Carried in a Staphylococcal Cassette Chromosome Genetic Element. Journal of Bacteriology 184, 3623–3629 (2002).

18. Chen, L. et al. Identification of a Novel Transposon (Tn6072) and a Truncated Staphylococcal Cassette Chromosome mec Element in Methicillin-Resistant Staphylococcus aureus ST239. Antimicrobial Agents and Chemotherapy 54, 3347–3354 (2010).

19. Lin, Y.-T. et al. A Novel Staphylococcal Cassette Chromosomal Element, SCCfusC, Carrying fusC and speG in Fusidic Acid-Resistant Methicillin-Resistant Staphylococcus aureus. Antimicrobial Agents and Chemotherapy 58, 1224–1227 (2014).

20. Kaya, H., et al. SCCmecFinder, a Web-Based Tool for Typing of Staphylococcal Cassette Chromosome mec in Staphylococcus aureus Using Whole-Genome Sequence Data. mSphere 3, 10.1128/msphere.00612-17 (2018).

21. O’Leary, N. A. et al. Exploring and retrieving sequence and metadata for species across the tree of life with NCBI Datasets. Sci Data 11, 732 (2024).

22. Kim, C. et al. Properties of a Novel PBP2A Protein Homolog from Staphylococcus aureus Strain LGA251 and Its Contribution to the β-Lactam-resistant Phenotype. J Biol Chem 287, 36854–36863 (2012).

23. Rolo, J. et al. Evidence for the evolutionary steps leading to mecA-mediated β-lactam resistance in staphylococci. PLOS Genetics 13, e1006674 (2017).

24. Madhaiyan, M., Wirth, J. S. & Saravanan, V. S. Phylogenomic analyses of the Staphylococcaceae family suggest the reclassification of five species within the genus Staphylococcus as heterotypic synonyms, the promotion of five subspecies to novel species, the taxonomic reassignment of five Staphylococcus species to Mammaliicoccus gen. nov., and the formal assignment of Nosocomiicoccus to the family Staphylococcaceae. International Journal of Systematic and Evolutionary Microbiology 70, 5926–5936 (2020).

